# A theoretical exploration of protocols for treating prosthetic joint infections with combinations of antibiotics and bacteriophage

**DOI:** 10.1101/2025.02.16.638499

**Authors:** Bruce R. Levin, Teresa Gil-Gil, Brandon A. Berryhill, Michael H. Woodworth

**Author notes:** Address correspondence to Bruce R. Levin, **Email:**. **Author Contributions:** Conceptualization: BRL, TGG, BAB, MHW Methodology: BRL, TGG, BAB Investigation: BRL, TGG, BAB Visualization: BRL, TGG, BAB Funding Acquisition: BRL Project Administration: BRL, TGG, BAB, MHW Supervision: BRL, TGG, BAB, MHW Writing– Initial Draft: BRL, TGG, BAB, MHW Writing– Review & Editing: BRL, TGG, BAB, MHW. **Competing Interest Statement:** The authors have no competing interests to declare.

## Abstract

With the increase in the placement of prosthetic joints and other hardware in the body has come an increase in associated infections. These infections are particularly difficult to treat due to the underlying bacteria generating matrices which resist clearance by immune system effectors or antibiotics. These matrices, biofilms, have two primary ways of being eradicated, either by physical removal by debridement or by killing the underlying bacteria. The viruses which kill bacteria, bacteriophage, are readily capable of entering into biofilms and eradicating the bacteria therein. Therefore, bacteriophage have therapeutic potential as a supplement to antibiotics for the treatment of prosthetic joint infections. In this investigation, we generate a mathematical-computer simulation model to explore the contributions of the innate immune system with antibiotics, bacteriophage, and the joint action thereof in the control of biofilm-associated infections. Our results question the proposition that bacteriophage are an effective addition in the treatment of prosthetic joint infections.

## Introduction

Over 1,460,000 knee and hip prosthetic joints will be placed in the US annually by 2050 [1, 2]. Prosthetic joint infections (PJIs) with *Staphylococcus spp*. and other bacteria are common sequela to these procedures and require additional surgery and extensive periods of antibiotic treatment to cure [1, 2]. A major limitation in treating PJIs is the formation of biofilms on the surfaces of the prosthetics, where bacteria are resistant to the action of antibiotics. Another limitation to treating PJI’s and other bacterial infections is the ascent of bacteria resistant to the treating antibiotics. There is a critical need to develop effective interventions to prevent and treat PJIs that can destroy the extracellular matrix, make the bacteria residing in biofilms more susceptible to antibiotics at physiologic concentrations in the joint space, and ultimately kill the underlying bacteria. Bacteriophage (phage) have many characteristics that suggest they could meet this need as a therapy that is complementary to antibiotic PJI treatment. In addition to their ability to kill and replicate on bacteria resistant to antibiotics, they penetrate biofilms and make these structures more amenable to antibiotic treatment [3-7].

In this study, we employ mathematical and computer-simulation models to design and optimize protocols for the joint action of antibiotics and phages in the treatment of PJIs. Our models identify the factors and key parameters contributing to the success and failure of antibiotics, phages, and their combinations in the treatment of PJIs. These parameters can be estimated experimentally and, most importantly, the predictions of the model can be tested with both in vitro and in vivo experiments. We discuss those experiments and their potential implications in the clinic for treating PJIs.

## Results

### Mathematical model of antibiotics and phage treatment of prosthetic joint infections

A diagram of the model we use to simulate the course of infections and their treatment with antibiotics and phage is presented in Figure 1.

**Figure 1.**
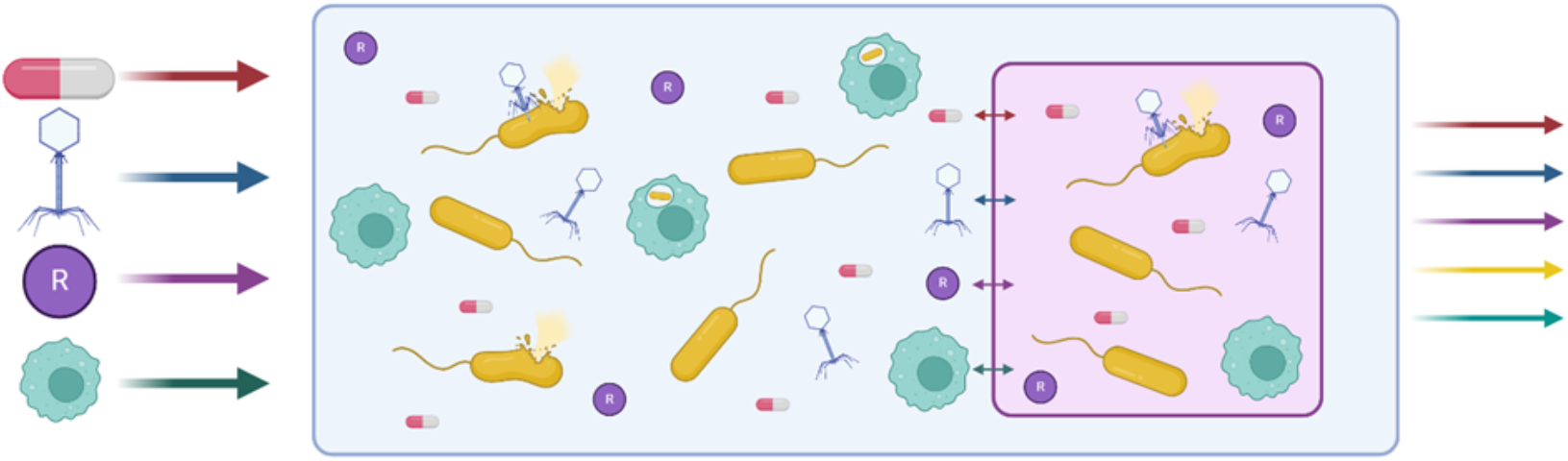
Diagram of a mathematical and computer-simulation model of prosthetic joint infections. The bacteria (yellow rods), bacteriophage (blue phage), resource (purple R particles), immune cells (green), and the antibiotic (striped pill) can be in two sites: the body, where they are planktonic (blue compartment), or in the biofilm (pink compartment). A limiting resource and immune cells enter the environment at a continuous rate. This is the same rate at which unconsumed resource, cells, phage, immune cells, and antibiotics exit the system. Antibiotics and bacteriophage can be dosed independently.

#### Pharmacodynamics of Antibiotic Treatment

We assume that resource (*R*, μg/mL) enters the environment at a constant rate and that the pharmacodynamics of the antibiotics and bacteria are modeled by a Hill function [8] where: ∏_*P*_*(A, R)* and ∏_*B*_*(A, R)* are the net growth/death rates of bacteria in the free, planktonic (*P*) and biofilm (*B*) habitats, respectively (Equation 1 and Equation 2). The parameters *v*_*maxP*_ and *v*_*maxB*_ (per cell per hour) are the maximum growth rates of the bacteria, and *v*_*minP*_ and *v*_*minB*_ (per cell per hour) the minimum rates of growth/maximum kill rates. *K* is the Hill coefficient and *k* (μg/mL) the Monod constant [9, 10]. To simulate the effect that the decreasing limiting resource concentration has on the physiological state of the bacteria, we include a term ψ(R) defined in Equation 3.

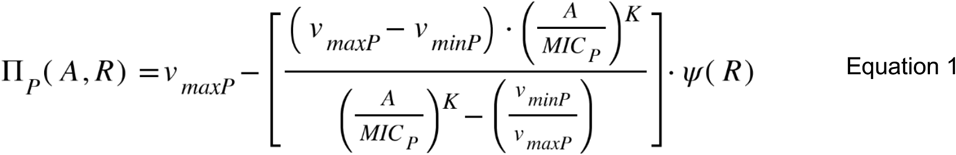

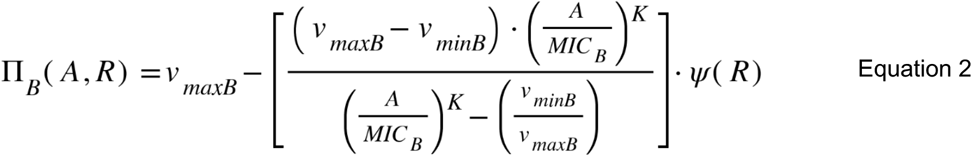

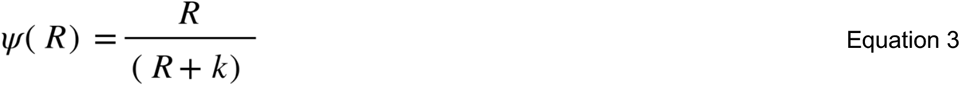

#### Mathematical Model of Phage and Antibiotic Treatment

To simulate the treatment of biofilm and non-biofilm bacteria with phage and antibiotics, we construct a series of coupled, ordered differential equations (Equations 4 through 11) given the following variables (Table 1), assumptions, and parameters (Table 2).

**Table 1.** Variables and their definitions.

| Variables | Definitions | Line Color in Figures |
| --- | --- | --- |
| R | The limiting resource | Green |
| A | The antibiotic | Black |
| I | Immune cells | Orange |
| IN | Immune cells containing bacteria | Red |
| NP | Free bacteria | Dark blue |
| VP | Free phage | Lilac |
| NB | Bacteria in the biofilm | Turquoise |
| VB | Phage in the biofilm | Dark pink |
| NAP | Free antibiotic-resistance bacteria | Light blue |
| NAB | Antibiotic-resistance bacteria in the biofilm | Light pink |

**Table 2.** Parameters, their definitions, dimensions, values, and their source.

| Parameters | Definition | Dimensions | Values | Source |
| --- | --- | --- | --- | --- |
| e | Substrate conversion efficiency | $\mu\text{g}$ per cell | $5\text{E}^{-7}$ | [13] |
| K | Hill coefficient |  | 1 | [8] |
| k | Monod constant | $\mu\text{g}/\text{mL}$ | 1 | [13] |
| $V_{\max P}$, $V_{\max B}$ | Maximum growth rates | Per cell per hr | 1 or 0.2 | |
| $V_{\min P}$, $V_{\min B}$ | Minimum growth/death rates | Per cell per hr | -5 or -0.5 | |
| $\text{MIC}_P$, $\text{MIC}_B$ | Minimum inhibitory concentrations | $\mu\text{g}/\text{mL}$ | 1 or 100 | |
| $A_{\text{MAX}}$ | Amount of antibiotic added at every dose | $\mu\text{g}/\text{mL}$ | 5 | |
| $\chi$ and $\gamma$ | Rate of flow into and out of the biofilm | Per hour | 0.001 | |
| dose | Dosing interval | Hours | 8 |  |
| C | Maximum resource concentration | $\mu\text{g}/\text{mL}$ | 1000 | |
| $w_P$, $w_B$ | Flow rate | Per hour | 0.1 | [14] |
| ip | Rate constant of phagocytosis | Per hour per mL | $1\text{E}^{-5}$ | [8] |
| $\beta_P$, $\beta_B$ | Burst sizes of phage | Particles per cell | 50 | [15] |
| $\delta_P$, $\delta_B$ | Adsorption rates of phage | Per hour per mL | $1\text{E}^{-8}$ | [15] |

We assume the phage adsorbs to the bacteria with rate constants and burst sizes, that depend on whether the bacteria are planktonic, *δ*_*P*_ and *β*_*P*_, or in the biofilm, *δ*_*B*_ and *β*_*B*_ (per hour per mL or particles per cell, respectively) [11, 12]. Antibiotics and phage are introduced into the planktonic habitat at defined doses and densities every *dose* hours and enter the biofilm from the planktonic habitat at a constant rate. The effective density of the antibiotics declines at a constant rate as does the density of the phage, but the latter may increase due to replication on the phage-sensitive bacteria. In addition to the bacteria that are sensitive to the action of the antibiotics and phage, we allow for sub-populations that are resistant to antibiotics. In our analysis of the properties of this model, we will consider situations where the antibiotic kills antibiotic-susceptible bacteria and the phage kills antibiotic-susceptible and antibiotic-resistant bacteria. Of particular interest in our analysis will be situations where the phage makes the bacteria in the biofilm more amenable to antibiotic treatment and elucidating the conditions under which resistance to the antibiotic will and will not ascend.

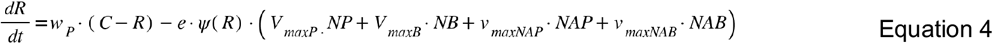

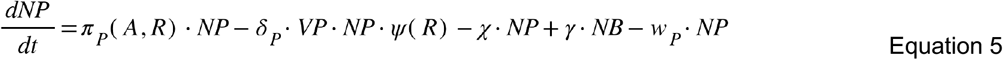

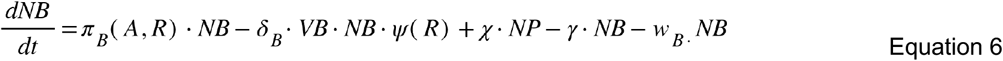

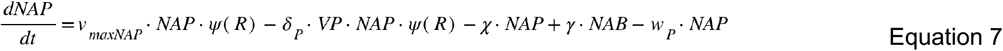

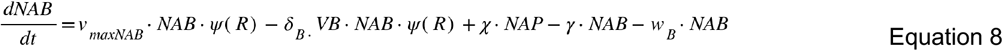

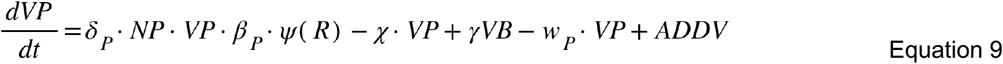

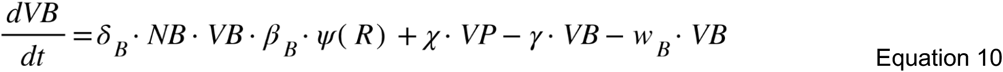

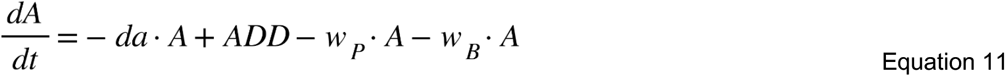

### Computer Simulations of the Treatment of Prosthetic Joint Infections with Antibiotics and Phage

In Figures 2, 3, and 4, we consider the changes in the densities of bacteria, phage, the immune cells, the concentration of the limiting resource, and the antibiotic concentration in the planktonic and biofilm states. The definitions of the variables and values of the parameters in the following simulations are those presented in Tables 1 and 2, unless otherwise stated.

**Figure 2.**
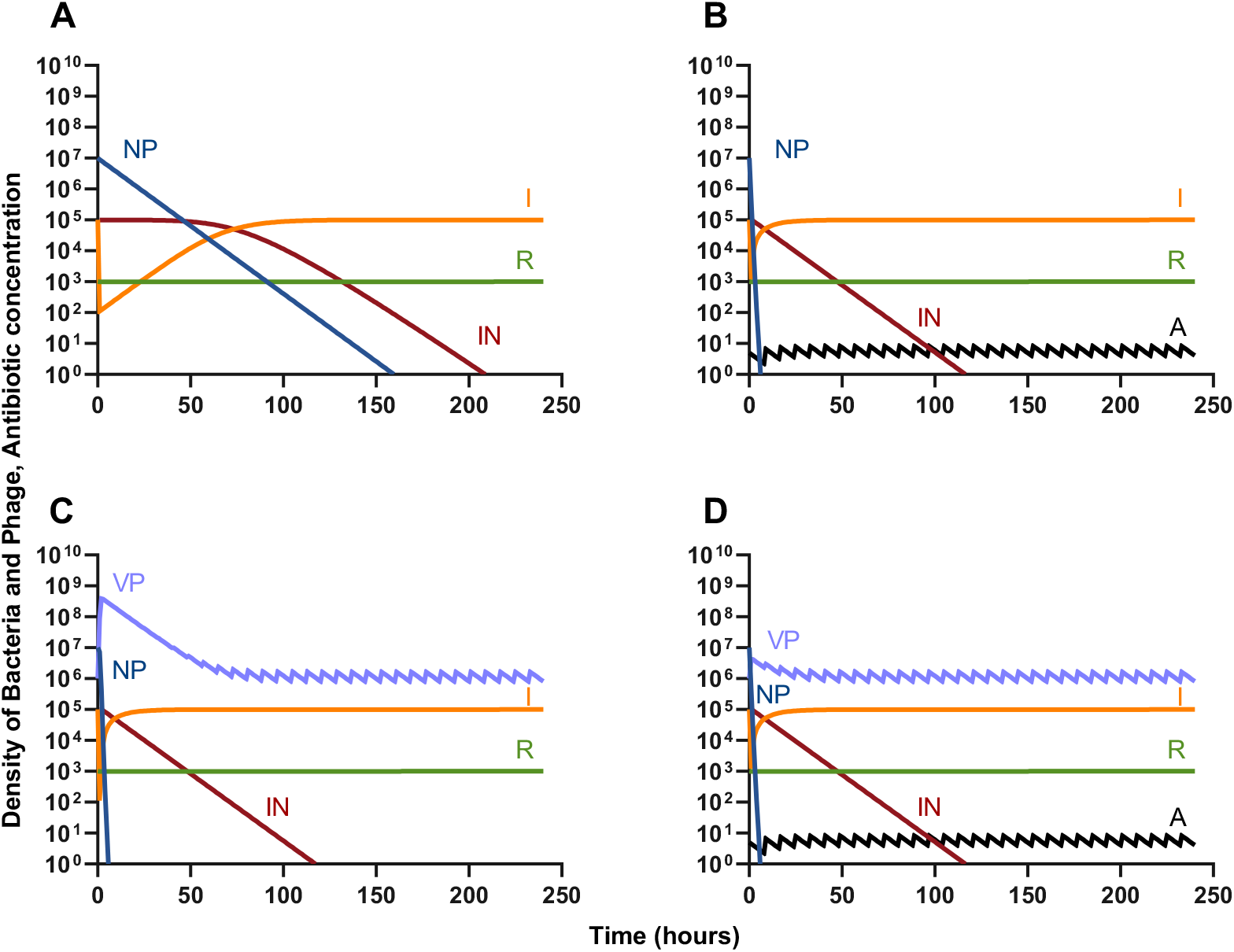
Simulated changes in the densities of bacteria, phage, and concentrations of the antibiotic and the limiting resource in the absence of a biofilm. A- An infection without treatment. B- Treatment with 5 μg/mL of an antibiotic. C- Treatment with 10^6^ PFU/mL of phage for an initial multiplicity of infection (MoI) of 0.1. D- Treatment with an antibiotic and phage.

**Figure 3.**
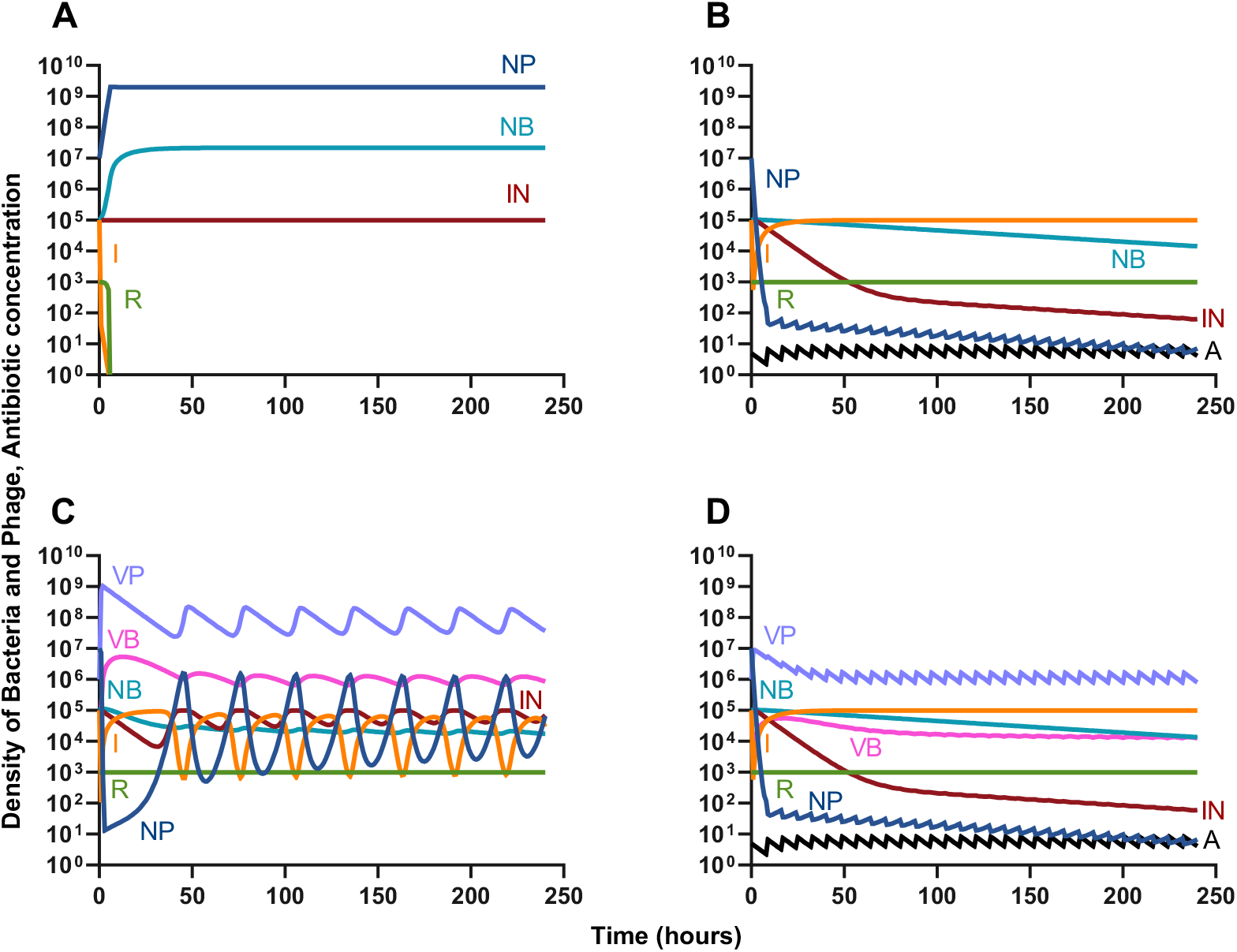
Changes in the densities of bacteria and phage in an infection with two habitats, planktonic and biofilm. A- An infection without treatment. B- Treatment with 5 μg/mL of an antibiotic. C- Treatment with 10^6^ PFU/mL of phage for an initial multiplicity of infection (MoI) of 0.1. D- Treatment with an antibiotic and phage.

**Figure 4.**
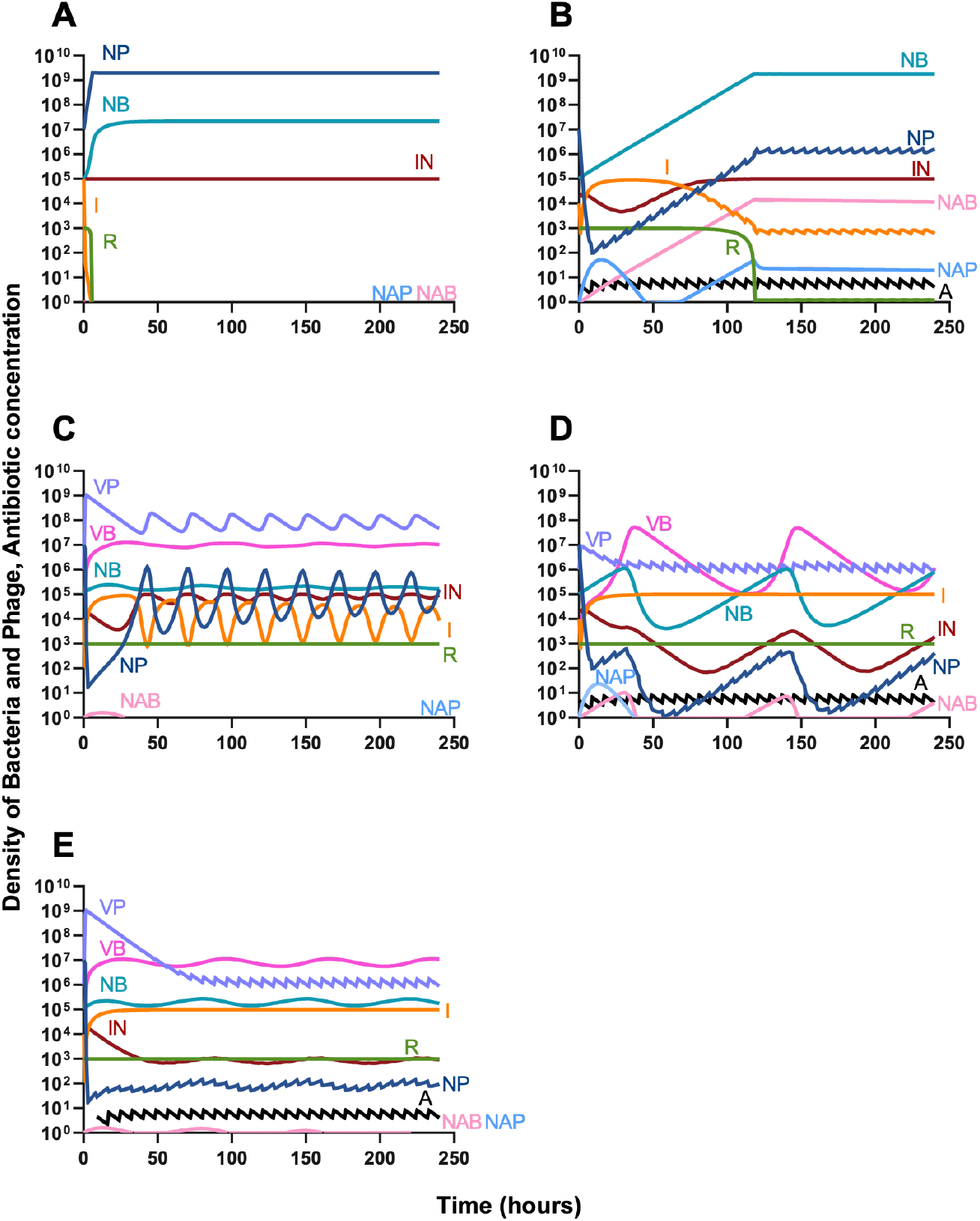
Changes in the densities of bacteria and phage in an infection with two habitats, planktonic and biofilm, and antibiotic-resistant sub-populations. A- An infection without treatment. B- Treatment with 5 μg/mL of an antibiotic. C- Treatment with 10^6^ PFU/mL of phage for an initial multiplicity of infection (MoI) of 0.1. D- Treatment with an antibiotic and phage. E- Treatment with a phage and then an antibiotic 8 hours later.

#### Treatment of Bacteria in the Absence of a Biofilm

In Figure 2 we follow the simulated dynamics in the absence of a biofilm site without the emergence of resistance. These simulations are initiated with 1/100^th^ of the densities of the populations and concentration of the limiting resource at equilibrium in the absence of treatment. In Figure 2A we consider the control of an infection by the innate immune system in the absence of treatment. In Figure 2B we consider the addition of fixed doses of a bactericidal antibiotic. In Figure 2C we consider the addition of fixed doses of bacteriophage. In Figure 2D we consider the joint action of an antibiotic and phage in controlling the infection. Notably, the time to clear the infection without any treatment is in excess of six days, while both individual treatments and the combined treatment speed up this clearance to less than seven hours. However, there are no major difference in the time to clearance between the three treatment conditions.

#### Treatment of Bacteria with a Biofilm without Antibiotic Resistance

In Figure 3 we follow these simulated dynamics with a biofilm site without the emergence of antibiotic resistance. These simulations are initiated with 1/100^th^ the densities of the populations and concentration of the limiting resource at equilibrium in the absence of treatment. In Figure 3A we consider the control of an infection by the innate immune system in the absence of treatment. In Figure 3B we consider the addition of fixed doses of a bactericidal antibiotic. In Figure 3C we consider the addition of fixed doses of bacteriophage which is able to enter to the biofilm. In Figure 3D we consider the joint action of an antibiotic and phage in controlling the infection. Unlike in the previous figure, the immune system alone is not able to control the infection, and, within short order, the system becomes resource limited. The addition of an antibiotic is able to suppress the planktonic cells, however it is unable to suppress the biofilm dwelling bacteria. The phage, on the other hand, are able to control the infection initially, however “boom-and-bust” type cycles emerge after the initial control of the planktonic cells [16]; much like in Figure 3B, the phage is unable to suppress the biofilm dwelling bacteria. Surprisingly, the combination of the antibiotic and phage yields a dynamic nearly identical to that of the antibiotic alone—implying that the bacteriophage is not affecting the dynamics. Taken together, these simulations predict that the action of the antibiotic will suppress that of the phage.

#### Treatment of Bacteria with a Biofilm with Antibiotic Resistance

In Figure 4 we follow these simulated dynamics with a biofilm site with the emergence of antibiotic-resistance. These simulations are initiated with 1/100^th^ the densities of the populations and concentration of the limiting resource at equilibrium in the absence of treatment. In Figure 4A we consider the control of an infection by the innate immune system in the absence of treatment. In Figure 4B we consider the addition of fixed doses of a bactericidal antibiotic. In Figure 4C we consider the addition of fixed doses of bacteriophage, which is able to enter to the biofilm. In Figure 4D we consider the joint action of an antibiotic and phage in controlling the infection. Like the above figures, the immune system is unable to control the infection, however the resistant sub-populations do not arise and, moreover, are lost. Treatment with just an antibiotic, is not able to control the infection; the antibiotic-resistant sub-populations are able to increase in density but not in relative frequency; and, the planktonic antibiotic-sensitive bacteria are maintained as a minor population due to transition from the biofilm associated antibiotic-sensitive cells. The addition of the phage, gives quantitively and qualitatively the same results as in Figure 3C as the antibiotic-resistant cells are rapidly lost. Interestingly, compared to 3D, the antibiotic in Figure 4D is not able to govern the dynamics of the infection as well, however the antibiotic-resistant sub-populations are still not able to ascend, but are maintained. Taken together, administration of a bacteriophage without antibiotic, is capable of eliminating any antibiotic-resistant sub-populations, which could make subsequent antibiotic treatment more effective; however, the co-administration of these antimicrobial agents is able to control but not clear the infection. When the antibiotic is delayed such that the phage is dosed first and then the antibiotic begins to be dosed 8 hours later, the cycles observed in Figure 4D are lower in magnitude and the antibiotic-resistant populations are lost (Figure 4E).

## DISCUSSION

Infections of prosthetic joints with *Staphylococcus spp*. and other bacteria are common sequelae to surgeries which place hardware into the body [17, 18]. These infections are particularly difficult to treat due to the emergence of biofilms on the hardware which decrease the efficacy of both the innate immune system and antibiotics [19]. Bacteriophage (phage), which are capable of penetrating the biofilm [3], would seem to be the perfect addition to antibiotic therapy to eliminate the bacteria which the antibiotics themselves are unable to kill. The results of the simulations we present in this study support the complimentary nature of phage in treating PJIs in only limited circumstances; particularly, when the phage is applied before the antibiotic. While generally, this may not be feasible in the clinic, in the case of PJIs it is, as these patients are generally not septic and these infections are usually subacute/chronic rather than acute and do not necessitate emergency treatment.

In the absence of a biofilm any sort of treatment—be it antibiotics, phage, or antibiotics and phage co-administered—speed up the time to clearance of the infection by the innate immune system. However, with a biofilm, both of these antimicrobial agents are able to suppress the infection but fail to clear it. Most notably, the antibiotic, and not the phage, seems to be the determining factor in the dynamics of treatment despite the inability to kill cells in the biofilm. The primary reason for the limited effectiveness of phage can be attributed to the predator-prey-type dynamics that emerge [16]. As the density of the bacteria is brought down by the phage, the rate of phage killing tends to zero as phage and bacteria interact in a density-dependent manner. Consequently, the phage begins to be washed out, their density declines, and thus the bacteria are able to increase in density again. On the other hand, antibiotics still kill at low bacterial densities, and thus the antibiotic does not cause the predator-prey-type cycles to emerge.

That being said, there is still likely clinical utility to phage. One is that these viruses can kill and control antibiotic-resistant bacteria and, as demonstrated above, could have utility if used before antibiotics to prevent the ascent of antibiotic-resistant sub-populations. Second, while the phage is not capable of clearing the infection, it does control the density of the infecting bacteria, potentially precluding the exhaustion of the innate immune system. Our conclusions are based solely on mathematical and computer-simulation models. It could well be that different results obtain when parallel experiments are performed. The utility of theory at large is in generating testable hypotheses and predictions which can be supported or rejected by experiments.

The models employed here do not take into account four things: i) the interactions which occur within a biofilm are in a structured environment rather than the mass-action environment simulated here [20]; ii) bacterial populations which are resistant to the phage could emerge; iii) populations which are resistant to both the phage and the antibiotic could also emerge; and iv) the role of the adaptive immune system. We do not believe these complexities require further modeling. First, models which consider structured habitats are much more complicated and their predictions more difficult to test. Second, there are many recently described phages for which resistance is extremely rare or cannot be generated [21, 22]. We believe the next step in the development of phage for treated PJIs is experimentation.

The starting conditions we use in our model can readily be recreated in vitro [23, 24]. For the in vitro experimentation many of the common biofilm models would successfully recreate the conditions simulated here. These experiments could also be performed in a range of model systems such as mice, as ultimately the utility of this work comes in the clinical application of phage and antibiotics to humans.

## MATERIALS AND METHODS

### Numerical Solutions (Simulations)

For our numerical analysis of the coupled, ordered differential equations presented (Equations 1-12) we used Berkeley Madonna [25] with the parameters presented in Table 2. Copies of the Berkeley Madonna programs used for these simulations are available at www.eclf.net.

## ACKNOWLEDGEMENTS

We thank the other members of the Levin Lab for their comments on an earlier version of this manuscript.

## Data Availability

The Berkeley Madonna programs used for these simulations are available at ECLF.net. All data are presented in this article or its supplementary materials.

